# Topokaryotyping demonstrates single cell variability and stress dependent variations in nuclear envelope associated domains

**DOI:** 10.1101/401539

**Authors:** Anamarija Jurisic, Chloe Robin, Pavel Tarlykov, Lee Siggens, Brigitte Schoell, Anna Jauch, Karl Ekwal, Claus Storgaard Sørensen, Marc Lipinski, Muhammad Shoaib, Vasily Ogryzko

**Author notes:** Present Address: [Anamarija Jurisic], Centre for Cancer Research and CellBiology, Queen’s University / Almac Discovery, BT9 7AE, Belfast, United Kingdom. [Chloe Robin], Centre de Biophysique Moléculaire CNRS UPR 4301, 45071 Orleans Cedex 2, France.

## Abstract

Analysis of large-scale interphase genome positioning with reference to a nuclear landmark has recently been studied using sequencing-based single cell approaches. However, these approaches are dependent upon technically challenging, time consuming and costly high throughput sequencing technologies, requiring specialized bioinformatics tools and expertise. Here, we propose a novel, affordable and robust microscopy-based single cell approach, termed Topokaryotyping, to analyze and reconstruct the interphase positioning of genomic loci relative to a given nuclear landmark, detectable as banding pattern on mitotic chromosomes. This is accomplished by proximity-dependent histone labeling, where biotin ligase BirA fused to nuclear envelope marker Emerin was coexpressed together with Biotin Acceptor Peptide (BAP)-histone fusion followed by (i) biotin labeling, (ii) generation of mitotic spreads, (iii) detection of the biotin label on mitotic chromosomes and (iv) their identification by karyotyping. Using Topokaryotyping, we identified both cooperativity and stochasticity in the positioning of emerin-associated chromatin domains in individual cells. Furthermore, the chromosome-banding pattern showed dynamic changes in emerin-associated domains upon physical and radiological stress. In summary, Topokaryotyping is a sensitive and reliable technique to quantitatively analyze spatial positioning of genomic regions interacting with a given nuclear landmark at the single cell level in various experimental conditions.

## INTRODUCTION

The mammalian genome is highly organized within the 3D space of the nucleus. This is exemplified by spatially defined localization of interphase chromosomes in nuclear sub-volumes and their non-random position relative to each other and to nuclear landmarks (1,2). The chromatin regions along the chromosomes are organized into domains through chromatin–chromatin interactions but also through their associations with the nuclear lamina, nuclear matrix and the nucleolus (3-5). Several studies demonstrated that disruption of the nuclear lamina leads to alterations in the organization of chromosomal domains (6,7) due to loss of contacts between the nuclear lamina and lamina proximal chromatin regions, termed lamina-associated domains (LADs). These domains are characterized by a lower gene density, transcriptional repression and are decorated with repressive chromatin marks (8). LADs contribute to the spatial chromosome organization by promoting proper chromatin folding during interphase (9). This is in line with the observations that upon transcriptional activation, certain developmentally induced genes relocalize from the transcriptionally repressive environment in the nuclear periphery towards nuclear interior (10-13). Changes in gene localization, can be also associated with different pathological states including laminopathies and cancer (6,14). Furthermore, altered spatial chromosome organization has also been shown to affect DNA replication and repair processes (15-18).

Over the last decade, significant insights into the higher-order spatial organization of eukaryotic genomes have been gained through advances in DNA imaging technology (19,20) and high-throughput biochemical techniques, such as Chromatin immunoprecipitation with massively parallel DNA sequencing (ChIP-seq) (21-24)], DNA methyltransferase identification (DamID) (25), or different versions of chromosome conformation capture; 3C, 4C, 5C and HiC (26-29). These approaches have a relatively high resolution, reaching from the nucleosome scale for ChIP-seq (i.e., 150 bp) and 1 kb for the latest version of chromosome conformation capture (HiC) (4). However, the main drawback of these approaches is the requirement for high numbers of cells, resulting in information averaged over a large cell population. Furthermore, the averaging might lead to the loss of critical information about correlations between the states of different genomic loci in individual cells. Another aspect taken into account is the heterogeneity, a fundamental property of cellular systems (30), which is lost while performing comprehensive chromatin conformation analysis. Finally, information regarding the proximity of genomic loci to a given nuclear landmark in various cell cycle phases is also lost using these methods.

Given that cell-to-cell variations may be genetic, epigenetic as well as stochastic in nature, understanding of the variability of intra-nuclear genome organization at the single cell level is extremely valuable for both fundamental and clinical research. An additional value of the single cell analysis is its potential to provide information about the correlations between different properties of individual cells including gene expression, variations in DNA sequence, cell cycle position, proteome or spatial chromatin organization, which is lost when the cells are analyzed in bulk. In this regard, several single-cell approaches have been developed in recent years to study different cellular states in individual cells, in particular, positioning of genomic loci with respect to each other and to nuclear landmarks (31,32). These approaches mainly based upon high throughput sequencing, require technically demanding and lengthy procedures while necessitate access to specialized equipment, in addition to being costly.

Here, we describe a novel and affordable methodology, Topokaryotyping, for quantitative analysis of spatial positioning of genomic regions interacting with a given nuclear landmark at single cell level. This robust microscopy-based approach relies on the fact that the information regarding spatial positioning of a particular genomic locus in the interphase nucleus is imprinted on mitotic chromosomes. Employing proximity utilizing biotinylation (PUB) (33) we biotin labeled histones in nuclear envelope (NE)-associated chromatin in interphase cells and detected biotinylation patterns on mitotic chromosomes. We demonstrate that the main advantage of this single-cell level approach is the dynamic mapping of NE-association events in individual cells. Our results indicate both cooperativity and stochasticity in the positioning of chromosome domains close to nuclear envelope. Moreover, we observed dynamic changes in the chromosome-banding pattern in NE-associated domains upon physical and radiological stress. Topokaryotyping is therefore, a simple approach, has relatively fewer steps than conventional methodologies with potential for increased throughput to study chromatin interactions to a fixed nuclear landmark.

## MATERIAL AND METHODS

### Recombinant DNA

To generate pCMV enhancer vectors, the pCDNA3.1 (Invitrogen) backbone was used. For the MoMuLV enhancer vectors, pMXs-IRES-Blasticidin or pMXs-IRES-Puromycin (Cell Biolabs) were used. The BirA ORF was PCR ampli?ed from the vector pBBHN (34). The BAP domain fragments were prepared by designing oligonucleotides and sequential PCR, followed by restriction digestion and insertion into the pCDNA3.1 or pMXs-IRES-Puromycin vector. The insert sequences were con?rmed by sequencing. The vectors generated were used to clone the ORFs Emerin, GFP, H2A, H2A.Z, H3.1 and macroH2A1.

### Cell culture

HEK 293T cells obtained from American Type Culture Collection were grown in Dulbecco’s modified Eagle’s medium with high glucose (PAA) and 10% dialyzed fetal bovine serum (FCS, Thermofisher). For transient transfection 4.10^5^ cells were transfected with 1μg PCDNA3.1 H3.1/H2A-BAP and 0.1μg PCDNA3.1 Emerin/GFP-BirA plasmid using a standard calcium phosphate precipitation method. Cells were analyzed two days after transfection. For the biotin labeling *in vivo*, biotin (Sigma) was added to the medium at final concentration of 5 μg/ml for 1h, while the pH was stabilized by addition of 50 mM HEPES (pH 7.35) to the medium. Hela S3 cells obtained from American Type Culture Collection were grown in Dulbecco’s modified Eagle’s medium with high glucose (PAA) and 10% dialyzed fetal bovine serum (FCS, Thermofisher). To generate HeLa S3 stable cell line expressing BirA-emerin and BAP-macroH2A1, BAP-H2A, BAP-H2A.Z or BAP-H3.1 the retroviral transduction was performed. 4×10^5^HEK 293T cells were transfected in a 6-well plate, with 0.5μg pCL-Eco packaging vector, 0.5μg pCMV-Gag-Pol plasmid and 1μg pMXs-IP BAP-macroH2A1, BAP-H2A, BAP-H2A.Z or BAP-H3.1 plasmids. Supernatants were collected 48h after transfection and used to transduce 4×10^5^HeLa S3 cells in presence of 4μg/mL of transduction enhancer polybrene (Sigma-Aldrich). After 7 days puromycin selection, HeLa S3 BAP-macroH2A1, BAP-H2A, BAP-H2A.Z or BAP-H3.1 expressing cells were second time transduced, as described above using instead pMXs-IB BirA-emerin or BirA-GFP plasmids followed by 7 days blasticidin selection.

### Cell treatments

HeLa S3 BirA-emerin and BAP-macroH2A1 cells grown in Dulbecco’s modified Eagle’s medium with high glucose (PAA) and 10% dialyzed fetal bovine serum (FCS, Thermofisher) were synchronized in G2/M phase in the overnight presence of CDK1 inhibitor RO-3306 (Sigma-Aldrich) at the final concentration of 5μM. Cells were exposed either to 5 Gray γ-irradiation, mild heat shock by incubation at 42°C for 2hrs/1h or 1h following 1h recovery at 37°C and biotin pulse labeled for 1h. Afterwards, mitotic spreads were prepared using methanol: acetic acid (as further described) and stained with streptavidin-Cy3 at 2μg/mL (Sigma S6402) and 4.6-diamidino-2phenylindole (DAPI, Sigma AF6057).

### Chromosome spreads preparation

Cells were synchronized and biotin pulse labeled as previously described. Biotin removal was followed by 45min treatment with colcemid (Sigma-Aldrich) at a final concentration of 0.1μg/ml. After mild trypsinization, mitotic cells were hypotonically swollen in preheated 75mM KCl at 37°C for 13 minutes. Cells were fixed by adding ice-cold fixative (3:1 methanol: acetic acid) following dropping onto slides. After drying, slides were then immersed at 37°C for 10 minutes in a permeabilization solution (120 mM KCl, 20 mM NaCl, 10 mM Tris pH 7.4, 0.5 mM EDTA, 0.1% Triton X-100, 0.1% Tween 20) and chromosomes were fixed for 20 min with 2% formaldehyde solution at 37°C.

### Cell Fixation and Immunostaining

2×10^5^HEK 293T cells were plated in serum free medium on 22×22mm coverslips, incubated for 30 min at 37°C with 5% CO_2_ to increase adherence and continued to grow in DMEM containing 10% dialyzed serum. HEK293T or HeLa cells were transfected as described earlier. After *in vivo* biotin labeling, cells were immediately washed with PBS, fixed with 4% formaldehyde for 20 min at room temperature (RT). Cells were permeabilized with 1x PBS containing 0,25% Triton X-100 for 10 min at RT. For immunostaining, fixed HEK cells or Hela cells were blocked 30 min in 3% PBS/BSA and incubated with streptavidin-Cy3 at 2μg/mL (Sigma S6402) or anti-polyHistidine at 1:4000 (Sigma H1029) and anti-emerin at 1:300 (ThermoScientific PA5-29731) which were detected respectively by an anti-mouse or an anti-rabbit conjugated to the Dylight 488 (ThermoScientific 35502 and 35552; used at 1:500 dilution). Immunolabeled samples were mounted and counterstained with Fluoroshield mounting medium containing the DNA intercalating dye DAPI (Sigma F6057). Cells were visualized with a high-resolution Spectral Confocal Leica SPE microscope, using a 63 x 1.4 oil immersion objective.

### Flourescence in situ hybridization (FISH)

Mitotic chromosomes spread and immobilized on slides were first incubated for 1h at RT in PBS/20% glycerol followed by four successive steps of freezing/thawing: samples were cryoprotected by soaking in PBS/20% glycerol and directly snap frozen in liquid nitrogen. After 3 washes in PBS, samples were incubated for 20 min at RT in a solution of 0,1M HCl following 2 new washes in a 2X SSC solution. Mitotic spreads were treated with 200μg/mL RNase in a 2X SSC solution for 1h at 37°C, washed 3 times in 2X SSC and equilibrated in 50% formamide/2X SSC for 12h at 4°C. Next, they were pre-incubated in an excess of 70% formamide/2X SSC for 10 min at RT before denaturation in the same solution for 5 min at 75°C. Hybridization mixture, containing the probe (Agilent Sure-FISH 19p13.11 CRFL1) in 49% formamide/EDTA, 300 ng/mL salmon sperm DNA in 2X SSC and 10% dextran sulfate was denatured 5 min at 75°C and hybridized to denatured metaphase spread for 3 days at 37°C. Samples were rinsed 5 min at 45°C in 50% formamide/2X SSC, followed by two washes of 5 min at RT in 2X SSC and 3 washes of 5 min in PBS. Mitotic spreads were counterstained with DAPI, covered with anti-fade solution, and stored at 4°C until analysis.

### Biochemistry and Western analysis

For Western blot analysis, cell nuclei were prepared by cell disruption in CSK buffer (100 mM NaCl, 300 mM Sucrose, 10 mM Tris pH 7.5, 3 mM MgCl_2_, 1 mM EGTA, 1.2 mM PMSF, 0.5% Triton X-100), and centrifugation for 10 min at 4000 rpm. Nuclei were resuspended in 1X NuPAGE LDS Sample buffer (Invitrogen, NP0007) with DTT (12.5 mM), sonicated, heated for 10 min at 72° C and loaded on 4-12% gradient Novex Tris-Glycine precast gels (Invitrogen, NP0315). After separation, the proteins were transferred to PVDF membranes and probed with primary antibodies against macroH2A1 at 1:1000 (Active Motif 39594) detected by an antirabbit-HRP antibody at 1:1000 (Roth 4750.1) according to the manufacturer’s protocoles. Expression of the BAP-tagged proteins was detected with an HRP-conjugated anti-polyHistidine antibody at 1:4000 (Sigma A7058) and biotinylated proteins were visualized with an HRP-conjugated streptavidin at 1:1000 (Sigma S2438). For streptavidin-HRP, an additional washing step with PBS/Tween 20 0.1% and NaCl 500 mM was performed.

### PUB-NChIP

Purification of total (non-biotinylated) and biotinylated chromatin from Hela S3 stably coexpressing BirA-emerin and BAP-macroH2A1 fusion proteins was performed as described elsewhere (35).

Briefly, HeLa S3 and HeLa BirA-emerin and BAP-macroH2A1 expressing cells were pulse labeled with 5 μg/ml biotin for 1 hour with the addition of 50 mM final HEPES added to stabilize pH. After the removal of the biotin, cells were washed with PBS following the washing in HB buffer (10 mM TrisHCl, pH 7.3, 10 mM KCl, 1.5 mM MgCl_2_, 0.2 mM PMSF, 10 mM β-mercaptoethanol). Next, cells were incubated in HB buffer on ice for 20 min and nuclei were released by breaking the cells with size B pestle in dounce homogenizer and spun at 800g for 15 min. Nuclei were resuspended in an equal amount of the TGME buffer (50% Glycerol, 0.5 M Tris pH 7.9, 1M MgCl_2_, 500 mM EDTA, pH 8.0 and 5 mM DTT). For micrococcal digestion, the nuclei were resuspended in 500 uL of TM2 buffer (10 mM Tris-HCl, 2 mM MgCl_2_, 0.1% Triton, 1 mM PMSF), 2.5 uL of 0.5 M CaCl_2_ and 25 uL of micrococcal nuclease (MNase) (0.2 U/uL; Sigma; N3755) were added before incubation at 37°C for 10 min. The reaction was stopped by adding 15 uL of 0.1 M EGTA. Nuclei were collected by spinning at 400g for 10 min at 4°C. After MNase digestion, the nuclei pellet was resuspended in 500 uL of prechilled 0.4 M salt extraction buffer (385 mM NaCl, 10 mM Tris-HCl [pH 7.4], 2 mM MgCl_2_, 2 mM EGTA, 0.1% Triton X-100, 1 mM PMSF). The tubes were rotated at 4°C for 30 min. Supernatant containing digested chromatin was separated from the remaining material by spinning at 400g for 10 min at 4°C. An equal volume of TM2 buffer was added to the extracted chromatin to arrive at a 0.2 M final salt concentration. The sample was centrifuged at maximum speed for 5 min at 4°C. For the preclearing, 300 μL of a suspension of Protein A agarose beads (Pierce 20334, Thermo Fisher) were washed three times with 0.2 M salt extraction. The bead suspension was combined with chromatin and rotated for 30 min at 4°C. The sample was centrifuged at maximum speed and supernatant transferred to a new tube. 300 μL of a suspension of sepharose-streptavidin beads (GE Healthcare; 17-5113-01) were washed three times with 0.2 M salt extraction buffer. The bead suspension was combined with the supernatant containing chromatin and rotated for 3 h at 4°C. Afterward, the beads were washed twice in 500 μL of 0.4 M salt extraction buffer containing Triton for 5 min at 4°C. DNA fraction was eluted twice by adding 100 μL of 1.5 M Ammonium bicarbonate and SpeedVac’ed. Afterwards, DNA was resuspended in 200 μL TE and an equal volume of phenol was added to sample and centrifuged for 5 min at maximum speed. Upper phase with DNA was transferred to a new tube. Two volumes of diethylether relative to the phenol phase were added to a lower protein containing aqueous phase and centrifuged for 5 min at maximal speed discarding upper phase. This step was repeated two times.

### Quantitative real-time PCR

2 μL of each sample was used per qPCR reaction. Samples were run in triplicate. The following mixture was prepared (one per sample): 7.5 μL qPCR master mix (Maxima SYBR Green/ROX qPCR Master Mix (2X) – ThermoFisher Scientific); 1 μL of forward and reverse primers mix (each at 10 μM); 4.5 μL ddH2O. The 2 μL from the DNA samples were then added and the PCR was performed. The list of primers is added as a table in the supplemental data. Following PCR program was used: 98°C, 30 s; then 40 cycles of 92°C, 1 s and 60°C, 15 s. Samples were run on Roche LightCycler 480 system.

### Next generation sequencing and analysis

For library construction, DNA fragments (100 ng) were converted to blunt ends by end repair using Ion Xpress Plus Fragment Library Kit (Thermo Fisher Scientific), purified with AMPure XP beads (Beckman Coulter) and ligated to Ion Xpress Barcode Adapters (Thermo Fisher Scientific). The resulting DNA fragments with a size of ∼200-220 bp (150-base-read libraries) were selected using E-Gel^®^ SizeSelect^™^ 2% (Thermo Fisher Scientific). The adapter-ligated library was then amplified by PCR using 8 cycles. DNA concentration was determined by qPCR and adjusted to 100 pM. The DNA fragments were templated on Ion Sphere Particles using the Ion OneTouch2 emulsion PCR instrument (Thermo Fisher Scientific) at a recommended molar concentration of 26 pM. Library sequencing was performed on Ion Torrent PGM system using Ion 318^™^ Chip Kit v2 BC and Ion PGM Hi-Q Sequencing Kit (Thermo Fisher Scientific). In addition to DNA from biotinylated chromatin, the sequencing was repeated using an input DNA (total digested chromatin) as a control. The sequencing data reported in this paper have been deposited in the Sequence Read Archive at the National Center for Biotechnology Information, (www.ncbi.nlm.nih.gov/sra, accession no. SRR6234698, SRR6234967 and SRR6235028). For data analysis reads were aligned to the human genome (hg19/GRCh 37).

HeLa LADs were obtained from GEO datasets (GSE57149). Downstream analysis and visualization was performed using Seqmonk analysis tool (https://www.bioinformatics.babraham.ac.uk/projects/seqmonk/). For visualization in genome browser images, the data were calculated as the log2 IP/input ratio across 10kb bins with smoothing correction applied for 5 adjacent probes. The scale for all genome browser images is log2 +/-2 with the exception of Figure 3D, which is log2 0-2. For Monte Carlo simulations, the genome was divided into larger 100kb fragments and regions overlapping LADs were tested against the genome in 10 000 simulations. HeLa RNA-seq data from the capped analysis of gene expression technique, as well as ChIP-seq data for Pol II, H3K4me3, H3K27ac and H3K36me3 were obtained from the ENCODE download portal (http://genome.ucsc.edu/encode/downloads.html).

### Quantitative analysis of EAD biotinylation patterns

The analysis consisted of taking 200 pictures of mitotic spreads per condition, selecting der19 chromosome from each spread based on its unique DAPI staining and finally annotating its biotinylation pattern. From the frequencies of various observed biotinylation patterns of the der19 chromosome we calculated the frequencies of each EAD. The biotinylation frequency of the EAD_C_ (constant) was 100% since no change in its biotinylation status was observed. Assuming independent associations, biotinylation frequency of EAD_V1_, EAD_V2_ and EAD_V3_, corresponds to the sum of all observed BPs where this domain has found to be biotin labeled as presented below.

**EAD_V1_** :BP_2_, BP_5_, BP_7_ and BP_8_ (0.2+0.08+0.02+0.06=0.36)

**EAD_V2_** :BP_3_, BP_5_, BP_6_ and BP_8_ (0.065+0.08+0.03+0.06=0.235)

**EAD_V3_** :BP_4_, BP_6_, BP_7_ and BP_8_ (0.03+0.03+0.02+0.06=0.14)

The calculation of expected frequencies was done as follows: on each BP, as previously mentioned, EAD_V1_, EAD_V2_ and EAD_V3_ can be either biotinylated (P-frequency of biotinylated EADs, calculated above) or non-biotinylated (Q-frequency of non-biotinylated EAD). Assuming independent associations,

**P+Q=1**

**EAD_C_**: P_0_=1, Q_0_=0

**EAD_V1_**: P_1_= 0.36, Q_1_=1-0.36=0.64,

**EAD_V2_**: P_2_= 0.235, Q_2_=1-0.235=0.765

**EAD_V3_**: P_3_= 0.14 and Q_3_=1-0.14=0.86.

Accordingly, the expected frequency for each BP can be calculated as follows:

BP_1_= Q_1_*Q_2_*Q_3_=0.42

BP_2_= P_1_*Q_2_*Q_3_=0.237

BP_3_= Q_1_*P_2_*Q_3_=0.129

BP_4_= Q_1_*Q_2_*P_3_=0.068

BP_5_= P_1_*P_2_*Q_3_=0.073

BP_6_= P_1_*Q_2_*P_3_=0.04

BP_7_= Q_1_*P_2_*P_3_=0.021

BP_8_= P_1_*P_2_*P_3_=0.012

Data were analyzed using statistical tests (GraphPad Prism 6 software) as explained in the Figure legends.

## RESULTS

### 2.1 Pulse and chase PUB allows for observation of NE-proximal genomic regions on mitotic chromosomes

We have previously published the Proximity Utilizing Biotinylation (PUB) method (33), which allowed specific labeling of a sub-fraction of the BAP-fused protein ‘X’ in proximity to a BirA-fused protein ‘Y’. In the current work, based on PUB principle (Figure 1A), we preferentially labeled LADs, utilizing Emerin, a constituent protein of inner nuclear membrane (36). First, in order to confirm its localization, we transiently expressed Emerin-GFP in HEK 293T cells. As shown in figure 1B, Emerin-GFP was detected predominantly at the NE. Once the peripheral localization of Emerin was confirmed, we next sought to achieve the preferential biotinylation of LADs by cotransfecting HEK 293T cells with Emerin fused to BirA and BAP fused to histones H2A or H3.1. Two days after transfection, cells were biotin pulse labeled for 1 h and subsequently fixed and stained with streptavidin-Cy3 and DAPI. The biotin signal was seen as a peripheral nuclear rim, consistent with BirA having specifically biotinylated BAP-histone (H2A or H3.1) fusions only in proximity to Emerin, localized at the NE (Figure 1C; Supplemental Figure 1A top). As evident from the images in Supplementary Figure 1B, in both BAP-H2A and BAP-H3.1 expressing conditions, majority of cells (60-70%) displayed a nice peripheral “ring shaped” biotinylation pattern (Supplementary Figure 1C). The cells that deviated from this pattern showed pan nuclear biotin signal, which may be due to high ectopic expression of the respective histones. Western blot analysis using anti-polyHis-HRP antibody confirmed the expression of BAP-H2A and BAP-H3.1 histones (Figure 1D, left panel, lane 1-4) regardless of their biotinylation status. Biotin signal detected with streptavidin-HRP showed that the BAP-H2A or BAP-H3.1 histone was biotinylated only when coexpressed with the BirA-fusion protein (Figure 1D, right panel, lane 3 and 4), in line with the results obtained using immunofluorescence (IF) microscopy (Figure 1C).

**Figure 1:**
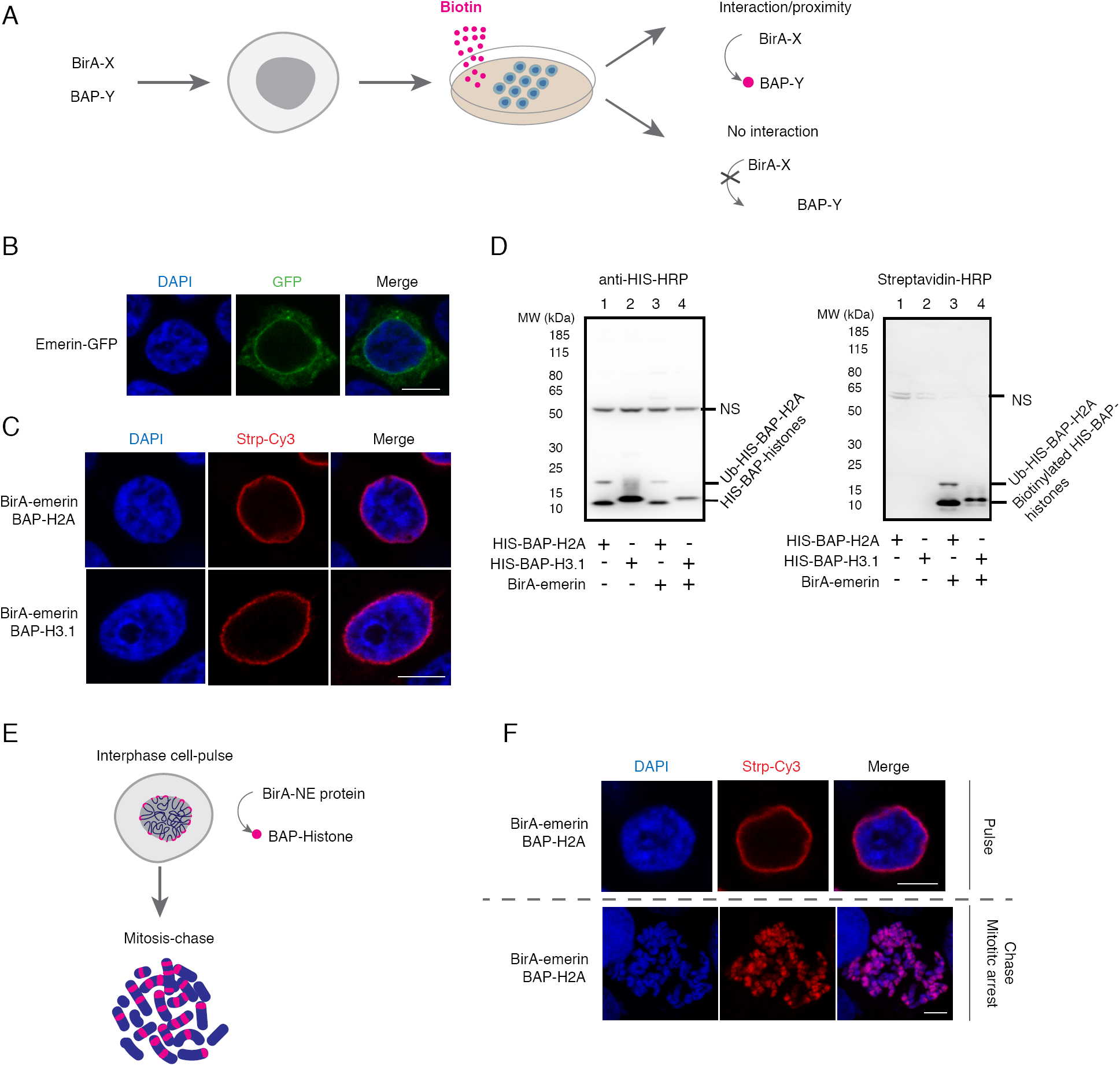
Proximity Utilizing Biotinylation (PUB) allows for detection of nuclear envelope-proximal regions on mitotic chromosomes. (A) The Proximity Utilizing Biotinylation principle. The PUB approach is based on coexpression of protein X fused to biotin ligase (BirA) and protein Y fused to biotin acceptor peptide (BAP). When the proteins are in sufficient proximity or interact with each other in cells harboring the corresponding plasmids, in the presence of biotin (pink), BirA specifically biotinylates the BAP peptide, resulting in a persistent label on the BAP-carrying protein. (B) Emerin is localized at the NE. 293T cells were transfected with emerin-GFP (green) and stained with DAPI (blue). Scale bar: 10 μm. (C) Biotinylation of NE proximal chromatin using BirA-emerin fusion. 293T cells were transfected with BirA-emerin and BAP-H2A or BAP-H3.1, followed by biotin pulse labeling and counterstaining with streptavidin-Cy3 (red) and DAPI (blue). Scale bar: 10 μm. (D) The biotin label depends on BAP-histone biotinylation. Western blot analysis was performed on nuclei from control (untransfected) and transfected 293T cells using anti-His-HRP (left) or streptavidin-HRP (right). The bands corresponding to BAP-histones, biotinylated BAP-histones, the ubiquitinated form of BAP-H2A (Ub-BAP-H2A) and non-specific signals (NS) are indicated. (E) PUB pulse and chase principle. BirA is fused to a nuclear envelope (NE) protein and BAP to a histone resulting in biotin labeling of chromatin proximal to the NE during interphase. The resulting biotin signals appear as bands on mitotic chromosomes. (F) PUB pulse and chase experiment. Interphase 293T cells coexpressing BirA-emerin and BAP-H2A are biotin pulse labeled followed by streptavidin-Cy3 (red) and DAPI (blue) staining (top). After 4h chase, biotin signal was revealed on mitotic chromosomes using the same staining dies as previously described (bottom). Scale bar: 10 μm.

As previously described (33), a unique feature of this approach lies in the covalent bond of biotin to the BAP domain, which persists even after the interaction with BirA has ended. This allowed us to monitor fate of the BAP-proteins at a specified moment after the biotinylation took place (Figure 1E). In particular, after pulse-labeling the NE-proximal chromatin in interphase, biotin was washed from the medium and the cells were allowed to remain in culture for another 4 hours. During this time, some of the cells had entered mitosis, carrying large chromatin domains revealed as biotinylated bands on mitotic chromosomes (Figure 1F). Since Emerin cannot be detected on the mitotic chromosomes, as the nuclear membrane has disintegrated, these biotinylated bands represent a record of the respective chromosome regions located close to the NE marker Emerin in the interphase nuclei. Confirming that the label is indeed Emerin-specific, the chromosome scale distribution of BAP histones was significantly more homogenous, as evidenced by the anti-polyHis-HRP staining (which detects BAP-histones regardless of their biotinylation status) of the same mitotic spreads (Supplemental 1A, bottom). Furthermore, the biotinylation status of BAP-histone distribution was also shown using BirA-GFP (uniformly distributed in the nucleus, Supplemental 1D, top) in PUB pulse and chase experiments (Supplemental 1D, top and bottom).

From this data, we conclude that PUB in pulse-chase format allows one to specifically mark the regions of mitotic chromosomes corresponding to the genomic regions that have been in contact with the NE during interphase.

### 2.2 Topokaryotyping, a PUB-based technique to detect interactions between chromatin and nuclear landmark as a staining pattern on mitotic chromosomes

The knowledge of the genomic identity of the biotinylated chromosome domains infers whether a given genomic region was associated with the NE in the interphase. This information is a useful aid in the reconstruction of the 3D genome topology, and moreover, given the work with individual spreads, it allows a single cell level analysis of genome organization in the preceding interphase. With this goal in mind, we have developed a four-step Topokaryotyping protocol which includes (i) PUB-labeling of chromatin in interphase cells using a NE protein fused to BirA and histone fused to BAP (ii) blocking cells in mitosis followed by mitotic spreads preparation, (iii) staining of mitotic chromosomes for biotin with the expected result being a partial staining of mitotic chromosomes, and iv) chromosome identification and quantitative analysis of relationship of chromatin-NE association events (Figure 2A).

**Figure 2:**
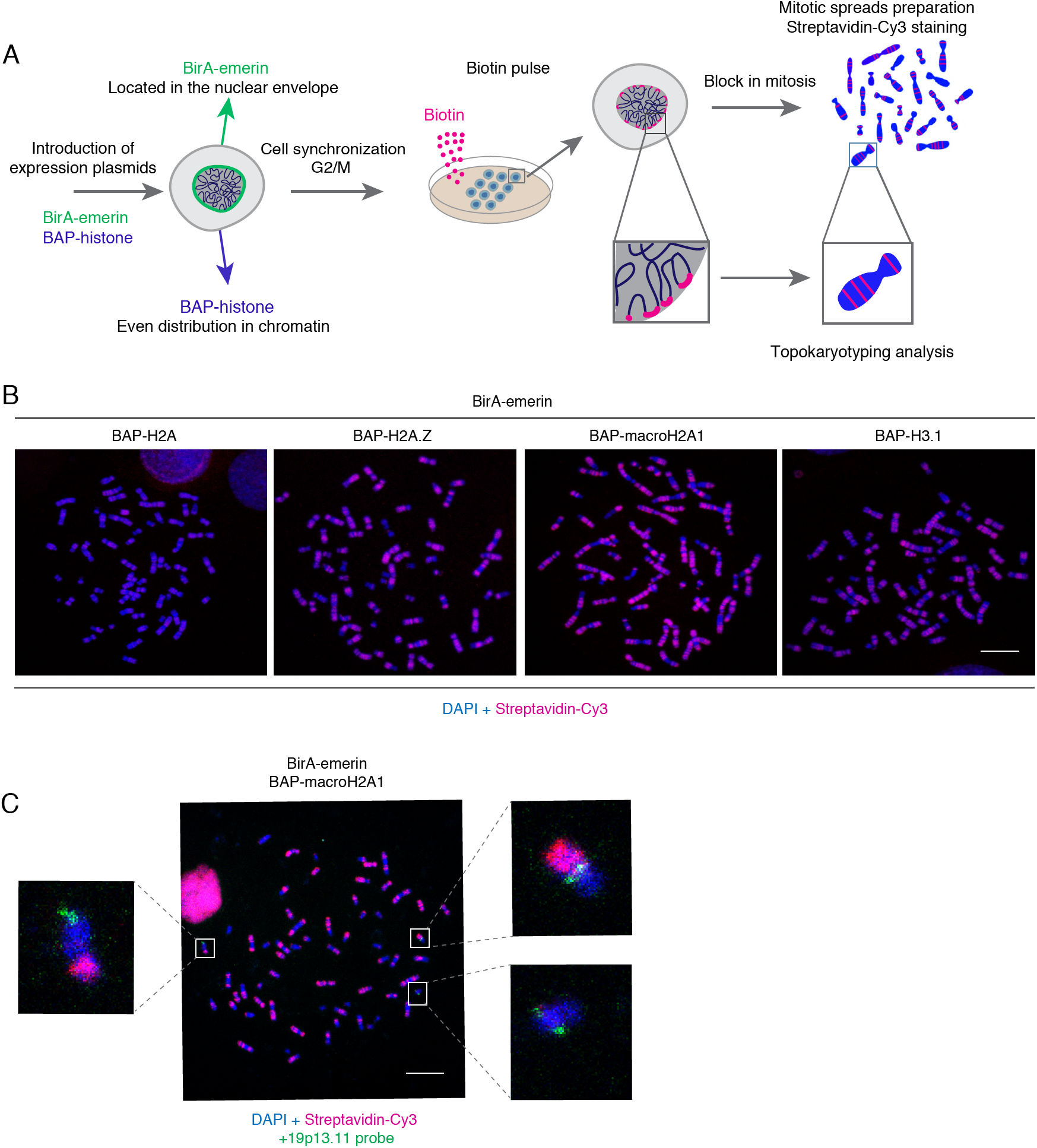
Topokaryotyping, a PUB-based technique to detect interactions between chromatin and nuclear landmark as a staining pattern on mitotic chromosomes. (A) Schematic representation of the principle for Topokaryotyping, based on labeling of chromatin in proximity with the NE. (i) BirA-fused to the NE-associated protein Emerin is expressed together with a BAP-histone fusion. (ii) After pulse-biotinylation, chromatin fragments that were close to BirA-emerin during interphase receive a biotin mark on their BAP-histones. (iii) Cells are blocked in mitosis and metaphase spreads prepared. Biotinylation is revealed in red and total DNA counterstained with DAPI. (iv) Chromosomal regions partially decorated in red can be further identified and analyzed. (B) Comparison of biotin signal on mitotic chromosomes between Hela S3 stable cell lines expressing BirA-emerin and BAP-H2A, BAP-H2A.Z, BAP-macroH2A1 or BAP-H3.1. Cells synchronized in G2/M phase during O/N treatment with RO3306 inhibitor were biotin-labeled and blocked in mitosis in the presence of colcemid. Afterwards, cells were hypotonically swollen and either permeabilized and fixed with methanol: acetic acid following dropping of mitotic chromosomes suspension onto glass slides. Spreads were stained with streptaviding-cy3 (red) and DAPI (blue). Scale bar: 10 μm. (C) PUB pulse and chase followed by Fluorescence *in situ* hybridization (FISH). Mitotic spreads from Hela S3 clone stably coexpressing BirA-emerin and BAP-macroH2A1 were prepared as described above and hybridized with a with a locus-specific probe for 19p13.11 (green) and counterstained with streptavidin-Cy3 (red) and DAPI (blue). Scale bar: 10 μm.

**Figure 3:**
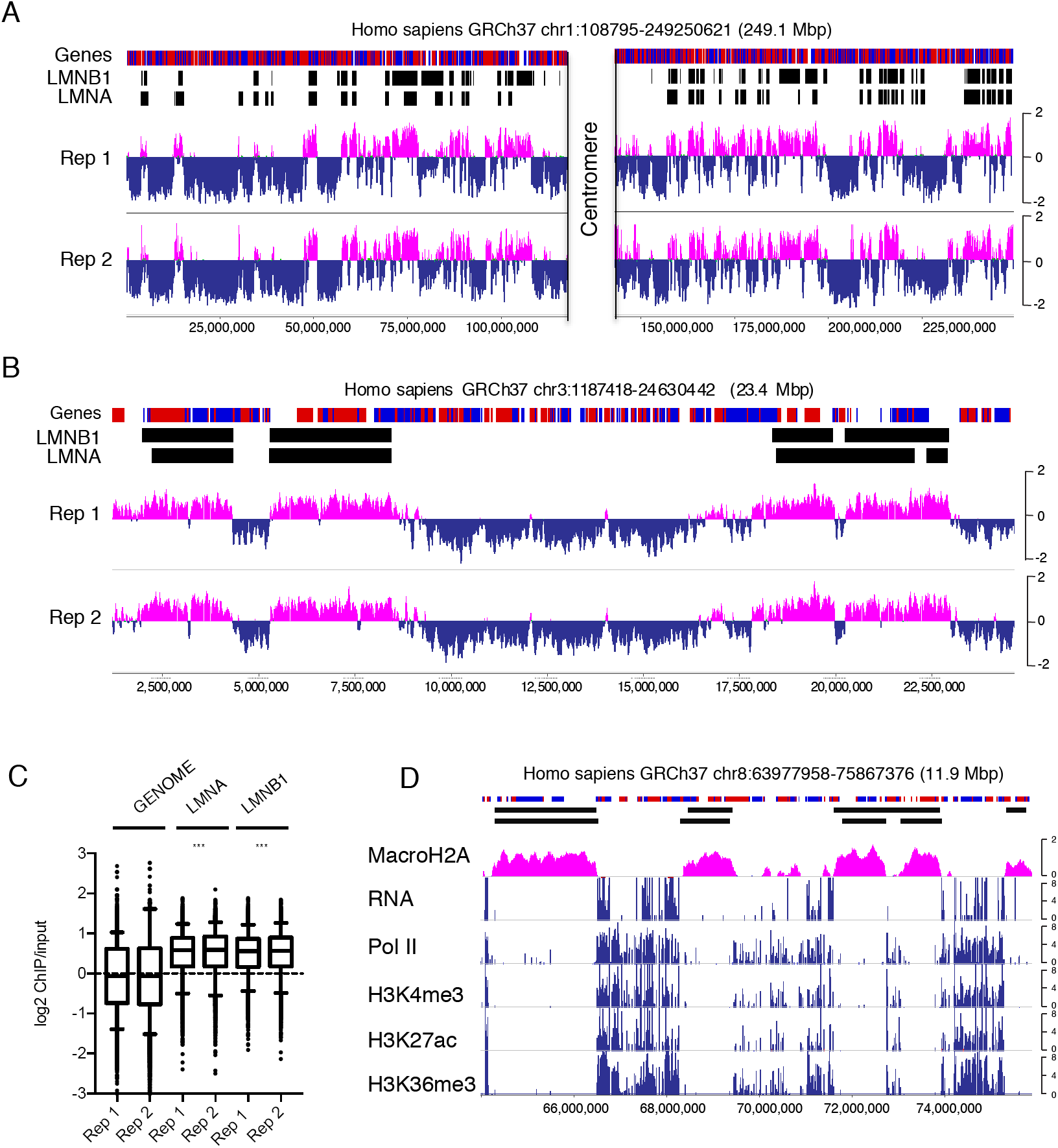
PUB-NChIP-sequencing shows that macroH2A1-emerin interaction occurs in lamina associated domains on a genome-wide scale. (A) Genome browser view of chromosome 1 showing LMNB1 and LMNA bound lamina-associated domains and the enrichment or depletion of PUB-NChIP signal relative to input. (B) Genome browser view demonstrating close correlation between enrichment and selected HeLa LADs on chromosome 3. (C) Boxplots showing enrichment of PUB-NChIP signal for BirA-emerin/BAP-macroH2A1 at LMNA and LMNB1 LADs relative the genome wide average across 100 kb regions. Statistical significance was determined from Monte Carlo simulations (**** p <0.0001). (D) In contrast to BAP-macroH2A1, RNA, Pol II occupancy and active histone modifications are depleted from LAD regions.

First, we aimed to verify whether the mitotic spread preparation was compatible with biotin detection. In order to obtain a consistent experimental model, we generated stable clonal HeLa cell lines coexpressing BirA-emerin and various BAP-histone fusions; BAP-H2A, BAP-H2A.Z, BAP-macroH2A1 or BAP-H3.1. (Supplemental Figure 2A and B). In order to increase the number of cells in mitosis, the clones were synchronized in G2/M phase with overnight (O/N) treatment using a CDK1 inhibitor (RO-3306), followed by a biotin pulse labeling for 1h. Cells were subsequently arrested in mitosis and subjected to mitotic spreads preparation. To our satisfaction, methanol-acetic acid technique was compatible with biotin detection (Figure 2B). In general, we observed biotinylation signal on mitotic chromosomes in case of all the BAP-histone variants used, with the strongest signal being detected when BAP-macroH2A1 was used together with BirA-emerin. This histone variant is known to be unevenly distributed at the nucleosome resolution level being more concentrated at the nuclear periphery (37,38). Furthermore, the biotinylation occurred only on the BAP-macroH2A1 fraction proximal to BirA-emerin, although BAP-macroH2A1 was distributed in a sufficiently uniform fashion at the chromosome scale, as shown by anti-polyHis-HRP signal (Supplemental Figure 3A). Moreover, western blot analysis of HeLa BirA-emerin and BAP-macroH2A1 expressing clone using an anti-macroH2A1 antibody showed that the expression levels of the endogenous and BAP-fused macroH2A1 were similar (Supplemental Figure 3B, lane 2 bottom and top band). Furthermore, comparison of the levels of endogenous macroH2A1 in untransduced control Hela S3 cells and in BirA-emerin and BAP-macroH2A1 expressing clone demonstrated constant expression levels in both samples (Supplemental Figure 3B, lanes 1 and 2, bottom band). Therefore, in order to study chromatin-NE associations using Topokaryotyping, for our further experimentation we decided to use the HeLa S3 clonal cell line stably coexpressing BirA-emerin and BAP-macro-H2A1. We did not detect any observable differences in cellular morphology and growth characteristics in cells stably coexpressing BirA-emerin and BAP-macro-H2A1, when grown in media with and without biotin. Although biotin has shown to be involved in several biological processes, including gene expression and cellular metabolism and cellular energy homeostasis (39,40), the molecular details of the role of biotin in these cellular functions remain elusive. However, to verify the cell cycle behaviour of HeLa cells (coexpressing BirA-emerin + BAP-macroH2A1) we analysed their cell cycle profile grown with and without biotin (Supplementary Figure 3C). The results showed no striking differences in the growth characteristics of HeLa cells grown in the presence or absence of biotin.

Next, we sought to determine the compatibility of PUB-based Topokaryotyping with fluorescence in-situ hybridization (FISH), which would facilitate the identification of individual chromosomes and the comparison of their biotinylation patterns across many spreads (Figure 2C). Therefore, for FISH labeling, HeLa clonal cell line (BirA-emerin/BAP-macroH2A1) was biotin pulse labeled and spread onto the glass slides using methanol-acetic acid technique. Afterwards, mitotic spreads were hybridized with probe for the 19p13.11 locus and stained with Cy3-streptavidin and DAPI. The biotin signal was detected on mitotic chromosomes together with the FISH signal (Figure 2C), demonstrating that Topokaryotyping protocol is compatible with the FISH procedure.

### 2.3 PUB-NChIP-sequencing shows that macroH2A1-emerin interaction occurs in lamina-associated domains on a genome-wide scale

Before embarking on the detailed analysis of chromatin-NE interactions at single cell level, we decided to verify that the biotinylated signal is indeed enriched in the chromatin domains positioned at the nuclear periphery. In this regard, we purified the biotinylated chromatin from HeLa S3 clonal cell line stably coexpressing BirA-emerin and BAP-macro-H2A1 using Proximity Utilizing Biotinylation native Chromatin Immunoprecipitation (PUB-NChIP) as described previously (35). Instead of analysing the protein fraction, we isolated DNA from the biotinylated chromatin pull down and performed high throughput sequencing in order to get insight into the genome wide distribution of the isolated DNA. Importantly, PUB-NChIP-seq of the DNA purified from HeLa S3 clonal cell line stably coexpressing BirA-emerin and BAP-macro-H2A1 showed high levels of enrichment at the LADs, which were identified previously in HeLa cells using anti-Lamin B1 and anti-Lamin A ChIP-seq (41). The enrichment of biotinylated BAP-macroH2A1 at LADs is visibly clear at both the chromosome level (Figure 3A) and at individual LADs (Figure 3B, Supplemental Figure S4A-D). To test if this enrichment was significant, the genome was divided into 100kb windows and Monte Carlo simulation using 10,000 iterations was performed to examine if probes overlapping LADs were significantly different from the genomic average. For both replicates, at both LMNA and LMNB1 LADs, the enrichment of biotinylated BAP-macroH2A1 observed was significant relative to the genome wide average (Figure 3C, p <0.0001). The same results were obtained at LADs determined in human fibroblasts by DamID (Supplemental figure 5A).

Furthermore, the vast majority of LADs obtained by Lamin B1 and Lamin A chIP-seq overlapping regions show positive PUB-NChIP-seq enrichment demonstrating the strong genome wide correlation between HeLa LADs and genomic sequences obtained by PUB-NChIP-seq. Supplemental figure 5B shows that the LADs overlapping genomic regions (magenta) and non-overlapping regions (blue) display a clear separation of signal. Consistently, the vast majority of LAD regions clearly showing positive enrichment.

Importantly, these results are consistent with the fact that BAP-macroH2A1 was preferentially biotinylated by BirA-emerin in the LADs, observed previously in the IF analysis (Supplemental Figure 2A). Furthermore, these data also indicate that the DNA isolated from biotinylated chromatin is compatible with high throughput sequencing and allows for identification of genomic regions proximal to BirA-fused protein of interest with sufficiently high resolution to match comparable well validated genome-wide techniques such as ChIP-seq.

Since LADs are generally considered to be principally inactive, gene poor regions (8), we further verified the validity of PUB-NChIP-seq data by analysing it against RNA-seq and ChIP-seq data from euchromatin marks found in transcriptionally active regions, such as H3K4me3 and H3K36me3 (42,43). As expected, we observed that LADs, while enriched in biotinylated BAP-macroH2A1, are depleted in RNA Pol II and the active histone marks H3K4me3, H3K27ac and H3K36me3 (Figure 3D, Supplemental figure 5C).

To assess the robustness of our approach, we compared biotinylation patterns at various genomic loci between different BAP-histone fusions using quantitative PCRs (qPCRs). In this regard, we performed PUB-NChIP followed by qPCRs in HeLa cells stably co-expressing BirA-emerin and BAP-H2A or BAP-H2AZ or BAP-macroH2A1. We designed qPCR primers for regions showing high enrichment in PUB-NChIP-seq data from different chromosomes (chr2, 3, 4, 5, 7, 10, 13, 15, 19) (Supplementary Figure S6). Our results indicated that cells expressing BAP-macroH2A1 consistently showed high enrichment at all the tested loci while BAP-H2A and BAP-H2AZ expressing cell lines showed slightly lower enrichment. These results support the notion that Topokaryotyping can be performed using other BAP-fused histones (canonical or variant). However, in case of detection and analysis of LADs, the assay works more robustly with BAP-macroH2A1, given its peripheral enrichment.

### 2.4 Analysis of der(19)t(3;19) chromosome reveals elements of cooperativity and stochasticity in the positioning of Emerin-associated domains

Next, to validate Topokaryotyping approach we conducted a single cell analysis of a model chromosome, der(19)t(3;19), present in HeLa cells (coexpressing BirA-emerin and BAP-macroH2A1). This single copy chromosome, further referred as der19, is originated via a translocation event (Supplemental 7A, 1) and easily identifiable by its morphology (Supplemental 7A, 2 and 3). Further characterization using partial chromosome painting (pcp) probes for the short (p) and long (q) arms of chromosomes 3 and 19 (Supplemental Figure 7B and C) demonstrated that the der19, contains material derived mainly from 3p and 19p (Supplemental Figure 7D). On a genome-wide scale chr19 has fewer NL contact sites, each with relatively low contact frequency as observed by Kind et al (figure 3 in Kind et al (32)). Consistent with this data, by PUB-NChIP-seq we see a very similar profile at chr19 (Supplemental Figure 8). However, the unique microscopic appearance of der19 facilitates the comparison of its biotinylation patterns across different spreads, which would be difficult otherwise, given that most chromosomes are present in more than one copy. Therefore, to reconstruct the NE-association map in single cells, the following analysis of der19 biotinylation patterns demonstrates a snapshot of information our approach could provide.

We compared the biotinylation pattern of der19 across 200 randomly selected individual mitotic spreads, prepared according to Topokaryotyping protocol (Figure 2A). Interestingly, we observed four stainable domains named “Emerin-Associated Domains” (EADs) (Figure 4A). One EAD at the tip of der19 was found biotinylated in all cases and was named EAD_C_ (for constant EAD, given this region is brought constantly in proximity with Emerin during interphase). Three other observed EADs, annotated as EAD_V1_, EAD_V2_ and EAD_V3_ (where, V stands for variable domains), were found to be associated with Emerin in a variable manner and accordingly, biotinylated in some but not all mitotic cells (Figure 4A). Altogether, der19 was found with eight possible biotinylation patterns (BP) composed from the various combinations of EADs (Figure 4B, 1), each reflecting a particular mode of association with the NE during interphase (Figure 4B, 2 and 3). The frequency with which each BP was observed was annotated as mean ± standard error from three independent experiments (Figure 4B, 4). Whereas, EAD_C_ was found stained in 100% of the mitoses, assuming independent associations, EAD_V1,_ EAD_V2_ and EAD_V3_ were found stained in 36%, 23.5% and 14% of the mitoses (corresponds to the sum of all observed BPs where this domain has found to be biotin labeled as calculated and presented in *Materials and Methods*). These observations indicate that the association with emerin during interphase occurs neither equally nor randomly at any part of the chromosome der19 (Figure 4C).

**Figure 4:**
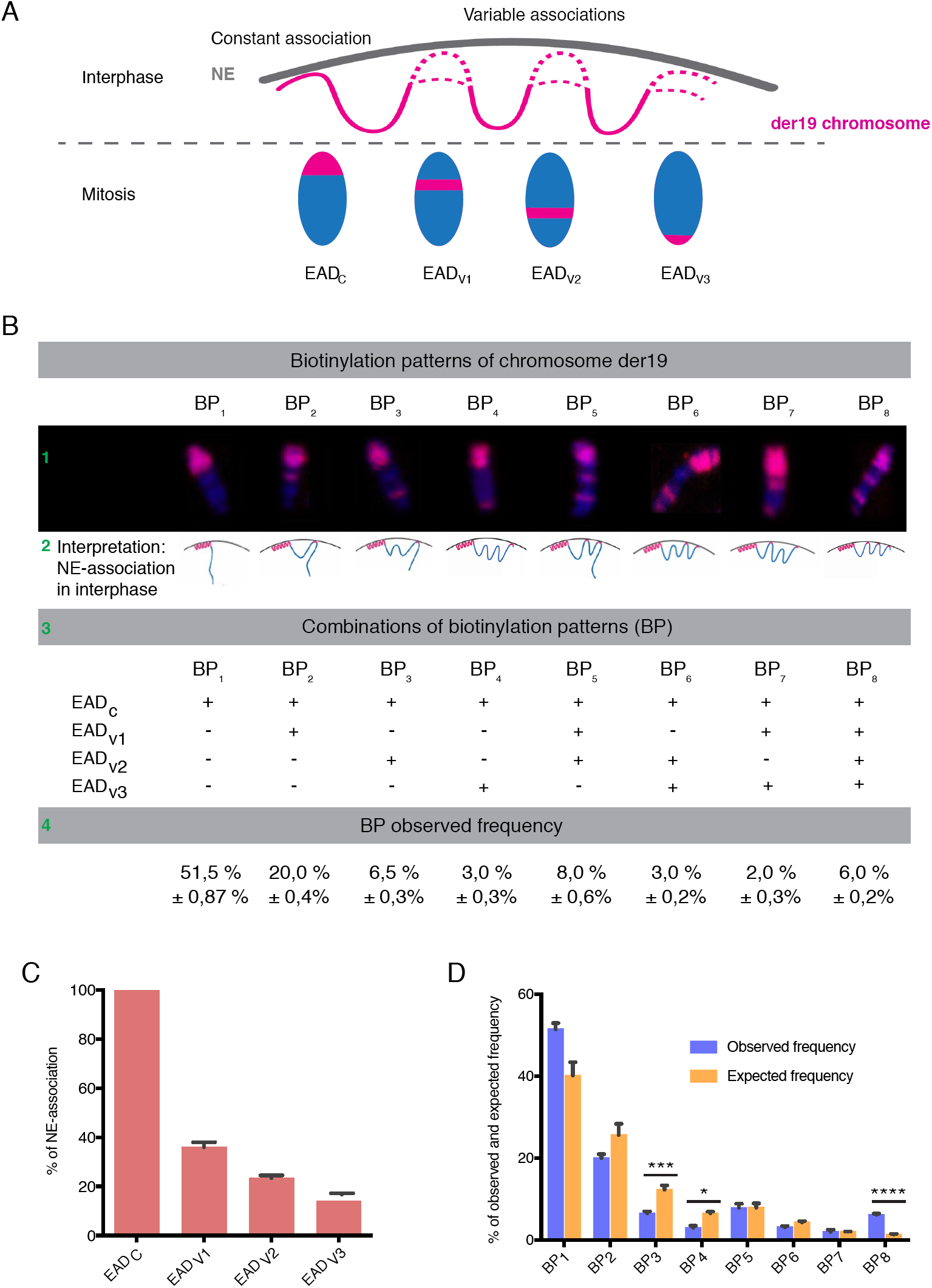
Topokaryotyping analysis of the der19chromosome reveals elements of cooperativity and stochasticity in the positioning of Emerin-associated domains. Mitotic spreads from Hela S3 clone stably coexpressing BirA-emerin and BAP-macroH2A1 were prepared using methanol-acetic acid protocol and stained with streptavidin-Cy3 (red) and DAPI (blue). (A) Schematic representation of emerin associated domains (EADs) in der19 chromosome. One domain is constantly associated with emerin (EAD_C_) while three other domains are associated in variable manner (EAD_V1_, EAD_V2_ and EAD_V3_). (B) Analysis of biotinylation patterns (BP) in der19 chromosome. Eight different BPs are identified (1), with their interphase NE-association interpretation (2), possible combination of EADs (3) and observed biotinylation frequencies (4). (C) Biotinylation frequency of EADs calculated from BP observed frequencies. (D) Comparison of observed and expected values of each BP where ** p <0.01; *** p <0.001; **** p <0.0001, determined by Chi-square test. Data are presented as means ± SEM of three independent experiments.

Next, we sought to determine the relationship of NE-association events among three variable EADs. The assumption was that if during interphase the association of each EAD with Emerin occurs independently from each other, then the biotinylation patterns were expected to be predictable from the individual frequencies observed for all EADs (detailed calculation is presented in *Materials and Methods*). The comparison of distribution between observed and expected frequencies for BPs (Figure 4D) reveals strong relationship among the NE-associations of the variable EADs (with the p-value describing the deviation from random distribution, corresponding to independent NE-associations). Assuming independence of the NE-associations, one should expect the pattern BP_3_ (only EAD_V2_ biotinylated) to be observed with the frequency 12.9%, whereas the pattern BP_4_ (the EAD_V3_ biotinylated only) should be observed with the frequency 6.8%. Instead, we observed 6.5 % and 3% for these patterns, respectively. The frequency of the pattern BP_8_ (EAD_V1,_ EAD_V2_ and EAD_V3_ biotinylated) deviated from the expected even more strikingly (6% vs 1.3%). For the rest of the BPs, we did not observe statistically significant difference between their expected and observed frequencies. This deviation from expected to observed frequencies strongly suggest that not all EADs associate independently. We discuss the implications of this observation in the discussion below.

### 2.5 Nuclear envelope-associations of der19 chromosome under physical and radiological stress show dynamic distribution

Once we established the biotinylation patterns of NE-association of der19 domains (EADs), using Topokaryotyping, we set out to investigate whether these association events are affected by various kinds of exogenous cellular stresses. To this end, the HeLa S3 clonal cell line (BirA-emerin and BAP-macro-H2A1) was synchronized in G2/M phase using a CDK1/2 inhibitor (RO-3306) for 16 h and further exposed to 2 h long mild heat shock at 42°C or 1 h heat shock together with an additional 1 h recovery at 37 °C. To induce radiological stress, cells were exposed to 5 Gy of ionizing radiation (IR). After the stress exposure, cells were biotin pulse labeled for 1 h, arrested in mitosis using colcemid and processed for mitotic spread preparation. This was followed by biotin detection and single cell analysis of der19, as described in Figure 4C and D. The schematics of the experiment are shown in Figure 5A and B (top). Before investigating the EADs’ frequency of association in these experimental conditions, we analyzed the cell cycle profile of cells synchronized in mitosis with and without IR. We did not detect any striking difference in the cell cycle profiles of untreated versus IR-treated cells (Supplemental Figure 9A). This is consistent with previous work (44,45) suggesting that high Polo-like kinase 1 activity accumulated in G2-arrested cells (CDK1 inhibitor) allows mitotic entry despite of remaining checkpoint signal or DNA damage.

To determine the effect of exogenous cellular stresses on NE-association of der19, we compared EADs’ frequency of association after HS and IR with that in control untreated cells (Figure 5A and B, bottom). In this regard, we found out that the NE-association of the variable EADs is highly sensitive to stress conditions. Intriguingly, the association of all three variable EADs with Emerin was decreased following heat shock, inducing an effect, which increased with duration of the shock (e.g., NE-association of EAD_V1_ for 1h HS was 23.2% vs 13.8% for 2h HS, as compared to 34% in control). However, this effect diminished when the cells were allowed to recover, showing restored distribution closer to the control one (e.g., NE-association of EAD_V1_ for 1h HS followed by 1h recovery was 27.5% vs 34% in a control, non-treated cells; Figure 5A, bottom). Following IR, the chromatin association of all variable EADs with emerin was also significantly decreased (e.g., NE-associations for EAD_V1_ after IR was 23.2% vs 34% in a control sample with the, Figure 5B bottom). Notably, we observed that the association frequency of constant EAD with Emerin was not affected by any stress, as it remained associated in 100% of analyzed mitotic spreads.

To further our investigation, we performed analysis of NE-association frequencies of BPs in each experimental condition. Given that BP_3_, BP_4_ and BP_8_ did not follow independent association trend (Figure 4D), in the following analysis we decided to focus on these three BPs. The comparison between observed and expected BP frequencies in all stress conditions showed that significant differences between these values for BP_3_ and BP_8_ are maintained when compared to control sample. BP_4_ follows the same trend as BP_3_ and BP_8_, however, the difference between observed and expected frequencies in this case is not statistically significant (Figure 5C-E, and Supplemental Figure 9B, C). These results indicate that NE-association of der19 with Emerin was decreased upon stress exposure, however, importantly, the overall relationship between associations of different variable EADs was not affected.

Finally, to confirm the changes observed in EADs shown in Figure 5B, we performed PUB-NChIP followed by qPCRs in HeLa cells co-expressing BirA-emerin and BAP-macroH2A1. We designed qPCR primers for genomic regions showing high enrichment in PUB-NChIP-seq dataset mainly from chromosome 3 and 19 (der19) but also from chromosomes 2 and 7. We compared the enrichment of these regions with and without IR treatment in a similar experimental setup as described in Fig. 5B. The results are consistent with our IF-based analysis where we saw no change in 3 loci from chromosome 3, which most likely belonged to constant EADs while there is a consistent reduction in 3 different loci from chromosome 19p (variable EADs), which as in Fig. 5B showed a reduction in Emerin binding after cellular treatment with IR. Apart from chromosome 19, genomic loci from chromosomes 2 and 7 also showed significant reduction after IR treatment, confirming that they belong to variable EADs.

**Figure 5:**
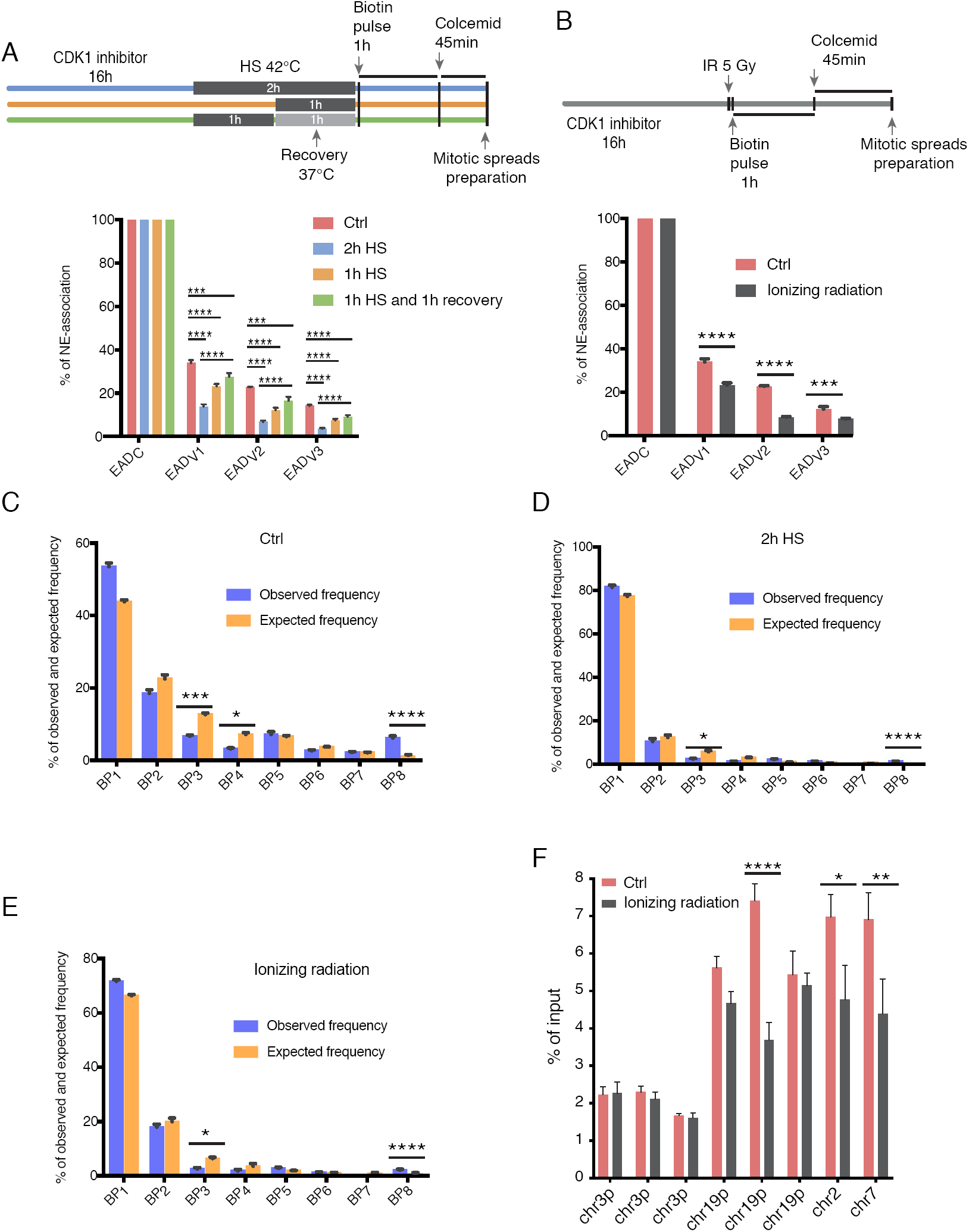
Nuclear envelope-associations in der19 chromosome under physical and radiological stress show dynamic distribution. (A) Top, experimental setup. Hela S3 clone stably coexpressing BirA-emerin and BAP-macroH2A1 was synchronized in G2/M phase in the presence of CDK1/2 inhibitor (RO3306) and exposed to heat shock at 42°C either 2h or 1h followed by 1h of recovery at 37°C. Afterwards cells were biotin pulse labeled for 1 h followed by biotin removal, cell arrest in mitosis and mitotic spreads preparation. Streptavidin-Cy3 and DAPI counterstained spreads were analyzed. Bottom, distribution of NE-association of EADs after heat stress. ** p <0.01; *** p <0.001, one-way ANOVA followed by Tukey’s post hoc test. All data are represented as means ± SEM from three independent experiments. (B) Top, experimental setup as in (A) where instead of heat shock, cells were treated with ionizing radiation (5 Gy). Bottom, distribution of NE-association of EADs after ionizing radiation. *** p <0.001; **** p <0.0001, one-way ANOVA followed by Tukey’s post hoc test. All data are represented as means ± SEM from three independent experiments. (C-E) Analysis of BPs after stress exposure (heat shock and ionizing radiation). Observed and expected frequencies of BP for individual cells grown in different experimental conditions are indicated. * p<0.05; *** p<0.001; **** p<0.0001, determined by Chi-square test. Data are presented as means ± SEM of three independent experiments. (F) Hela S3 clone stably coexpressing BirA-emerin and BAP-macroH2A1 was synchronized and treated as in (B). PUB-NChIP was performed as described earlier followed by qPCRs. qPCR primers for genomic regions showing high peaks in PUB-NChIP-seq data mainly from chromosome 3 and 19 but also from chromosomes 2 and 7 as well. The bars represent % of input in each sample ± SEM from 3 independent experiments. * p<0.05; ** p<0.01; **** p<0.0001 determined by 2-way ANOVA and Sidak’s multiple comparison tests.

## 3. Discussion

We describe here a novel and highly sensitive microscopy-based methodology termed Topokaryotyping, to analyze the 3D genome topology in individual cells with chromosome band resolution. It provides both topological and karyotypic information as illustrated by an example of the dynamic mapping of chromatin-NE association events of a unique chromosome, der19. This is analogous to the top-down analysis in proteomics (46), which can detect correlations between modifications of different amino acid residues in individual protein molecules, the information that is mostly lost when the proteins are fragmented by trypsin. Similarly, the chromatin analyses that involve DNA fragmentation from a population of cells (including ChIP or other biochemistry-oriented approaches) lose information about the correlations between the chromosome regions located far from each other.

This approach provides an unprecedented opportunity to observe, with high statistical significance, the relationship of NE-association events at the individual cell level. Focusing on these observations, we have obtained a glimpse at what kind of mechanistic insight(s) into the 3D nuclear organization our approach can provide. For example, we identified constant (EAD_C_) and variable emerin-associated domains (EAD_V_) in genetically identical HeLa cells, which is consistent with previous observations of cell-to-cell variability in NE-associations (32,47), and therefore, can be considered as their independent confirmation. The frequencies of association of these individual domains of der19 chromosome evidently differ from each other (Figure. 3C). Many alternative models can explain these differences, the most relevant being (i) ‘Independence and affinity gradient’ where, the both EAD_C_ and EAD_V_ domains associate with NE independent from each other, but have different propensity for it. One should expect all possible combinations of NE-association events, with the expected probability to detect each pattern computable from the independence assumption, (ii) ‘Cooperativity’, or ‘NE-association spreading’ where, EAD_C_ serves as the chromosome der19 anchor to NE, and the NE-association of each EAD_V_ domain requires the NE-association of the previous domain(s). Only a subset of patterns is expected in this case. This model is similar to the spreading of heterochromatin studied on Position Effect Variegation and silencing in experimental systems (48) and is easy to understand mechanistically, (iii) ‘Mutually exclusive NE-association and affinity gradient’ where, opposite to the model (ii), the NE-associations of each EAD_V_ domain is mutually exclusive, e.g., due to mechanical constraints. A different subset of patterns should be observed in this case. However, our results demonstrate a pattern distribution of emerin-associated domains, which strongly argues against the latter possibility. These observations suggest that the rules governing the NE-association of der19 favor a model between the first two extremes described above. Given strong relationship between the three EAD_V_, there is an element of cooperativity, but also, as manifested by patterns with independent NE-associations, that of stochasticity, the latter feature having many precedents in single cell level analysis (49). Moreover, the parallel (and a possible link) with heterochromatin spreading also raises a question whether the current picture of the latter phenomenon is oversimplified, and underscores the potential value of single cell level analysis in this case as well.

The unique feature of our approach is the ability to readily generate hundreds of spreads and subsequently detect NE-association events for many loci in individual cells. If combined with automated image acquisition and recognition, our approach provides a new way for preliminary screening for biological activities addressing different aspects of nuclear organization in different experimental conditions (i.e., separately evaluating general propensity of a chromosome domain for the NE-association and the cooperativity in NE-association between different domains). In this respect, the data on the stress response are intriguing. Consistently with the previous studies reporting the heat-induced changes in NE (50), we observed that treatment with heat shock decreases NE-association of der19 chromosome domains in a reversible dose-dependent manner. It has been reported that the heat shock proteins are induced in response to elevated temperature (51) together with the increased association of nuclear proteins with the nuclear matrix (52). It is therefore, attractive to speculate that denaturation and aggregation of nuclear proteins affects the physical properties of chromatin, possibly affecting chromatin-NE associations. Furthermore, the observed decrease in chromatin-NE association in IR treated cells could be explained by changes in irradiation-induced chromatin dynamics. Accordingly, upon induction of double strand breaks, chromatin decondensation has been observed at damaged sites (53,54), as well as, enhanced mobility of damaged chromatin in eukaryotic cells (55,56), which in turn affects chromatin-NE association. Importantly, affecting the NE-association of variable EADs in der19 chromosome, overall there was no statistically significant effect on the NE-association cooperativity in the case of these two stresses (Figure 5C-E, and Supplemental 9B, C).

The observation of multiple NE-associated domains on each chromosome is consistent with the well-known notion of LADs. Accordingly, the sequencing of the biotinylated chromatin from the HeLa cells used in this study (PUB-NChIP-seq), reveals strong overlap with the known LAD localizations previously identified by ChIP-seq analysis for Lamin A and Lamin B1 proteins (41,57). Furthermore, our PUB-NChIP-seq data from HeLa cells coexpressing BirA-emerin and BAP-macroH2A1 overlaps with LADs identified by DamID in human Tig3 cells (8,58). Also, as expected, this overlap is not perfect, most likely due to the presence of so-called cell-type specific LADs (47). In addition, comparing across cell types makes it impossible to accurately correlate various techniques, as it is not possible to know to what extent stronger or weaker correlations would reflect – since any differences could be related to cell type or the method used.

The importance of single cell based approaches lead to further adaptation of commonly used methods to study spatial chromosome organization. For instance, DamID based on single cell DNA sequencing by Kind et al (32) is conceptually similar to our technique (we fuse the biotin-ligase BirA to the protein of interest and, rather than DNA, we label a histone). On the other hand, Kind et al used the haploid KBM7 myleoid leukaemia cell line, with a distinct genetic profile that differs from HeLa cells making any direct comparisons between the two data sets challenging. However, our results are consistent with Kind *et al*, with respect to relatively low contact frequency of chr19 (Supplemental Figure 8). The weak nuclear lamina association of chr19 observed with genomic techniques such as DamID-seq, Lamin ChIP-seq and now PUB-NChIP-seq would appear to be conserved between the two cell types KBM7 and HeLa. Despite of fewer NL contact sites of chr19 observed in genome-wide sequencing data, we see a consistent der19 nuclear lamina attachment by topokaryotyping, which may relate to telomeric sequences that cannot be effectively mapped and/or the extra-chr19 sequences that become incorporated into chr19 in HeLa cells which have significant re-arrangements owing to a process known as chromothrypsis (59). Though it is difficult to draw conclusions between single cell DamID in one cell line and topokaryotyping in another, we would suggest that the nuclear lamina attachment observed with the der19 chromosome found in HeLa may come from the re-arranged portion that maps to chr3 or from telomeric repeats in chr19 which are not mappable due to their repetitive nature. While this is not the focus of our study it highlights the usefulness of the topokaryotyping approach providing additional information that is challenging to obtain through other genome-wide mapping approaches.

Our results are mostly in agreement in respect to the observations of cell-to-cell variability and cooperativity of the NE-association events. We would like to emphasize that the ease of obtaining positioning information with our approach allows one a relatively quick analysis of different experimental conditions. Moreover, our approach is free of one of the major problems encountered in sequencing-based approach, i.e., the interpretation of NE-association of repetitive sequences (60). However, there may be some technical challenges that may be time consuming while setting up the assay. These include generation of clonal stable cell lines coexpressing the BirA and BAP-fusion proteins of interest and validate the expression levels of the fusion proteins. Furthermore, one of the main drawbacks of Topokaryotyping is the lower resolution in comparison to other techniques such as ChIP-seq or DamID-seq. Nonetheless, the ability to easily perform and analyse Topokaryotyping experiments increases its versatility and potential to use.

Topokaryotyping imposes as a robust technique that could provide preliminary data on the effect of chromatin composition on various aspects of 3D genome organization. For instance, using histone mutants as BAP-fusion proteins (e.g. mutation in lysine 9 or lysine 27 in histone H3) one can study the role of individual histone residues (or their posttranslational modifications) in chromatin-lamina interactions. In this regard, comparison of NE-association events in wild type versus mutant BAP-histones provides clues to the role of various histone residues (PTMs) in determining the chromatin-NE contacts on different chromosomes in various experimental conditions. Furthermore, use of histone variants as BAP-fusion proteins or their mutated counterparts can add another dimension to this analysis. Alternatively, using RNA interference or CRISPR-based knockouts for different chromatin organizing proteins (such as CTCF, cohesion, condensins, heterochromatin protein 1, Lamins), one can study their role in NE-association on different chromosomes at single cell level. Owing to the precise timing of biotinylation pulse, a cell cycle-based analysis can also be done. Another interesting application of Topokaryotyping can be a preliminary assessment of the role of structural variants of genes identified in different cancers in their ability to disrupt chromatin-NE associations. This kind of analysis can be performed coupled with FISH probes against that particular variant.

Notably, the scope of our approach is not limited to the NE-association, rather it can be implemented to any fixed nuclear landmarks such as nucleolus, Cajal bodies or PML bodies. Importantly, however, the protein marker fused to BirA should be associated sufficiently strongly to the nuclear domain of interest, to decrease unspecific chromatin biotinylation due to the mobility of BirA-fusion protein. Accordingly, we expect that the proposed methodology will extend the toolbox of the available methods to analyze nuclear organization at the single cell level and will find numerous other applications in cellular biology, in particular, epigenetics and chromatin biology.

## DATA AVAILABILITY

PUB-NChIP-seq raw and processed data have been deposited in the NCBI gene expression omnibus (GEO) under accession number GSE118996. Also, the sequencing data reported in this paper have been deposited in the Sequence Read Archive at the National Center for Biotechnology Information, (www.ncbi.nlm.nih.gov/sra, accession no. SRR6234698, SRR6234967 and SRR6235028). For data analysis reads were aligned to the human genome (hg19/GRCh 37). HeLa LADs were obtained from GEO datasets (GSE57149). HeLa RNA-seq data from the capped analysis of gene expression technique, as well as ChIP-seq data for Pol II, H3K4me3, H3K27ac and H3K36me3 were obtained from the ENCODE download portal (http://genome.ucsc.edu/encode/downloads.html)

## SUPPLEMENTARY DATA

Supplementary Data are available at NAR online.

## ACKNOWLEDGEMENT

We are grateful to Dr. Vassilis Doucas for his constructive discussions in the early stages of the project and Ms. Sophie Salome-Desnoulez for assistance in confocal microscopy. We thank Dr. Alain Bernheim for his valuable advices in cytogenetics and Dr. Joelle Wiels for reading the manuscript. We thank Ms. Daria Marinovic for assistance in artwork for the figures. We would also like to thank Dr. Undine Mechold for useful discussions. We state and sincerely lament that Dr. Vasily Ogryzko passed away during the course of the study.

## AUTHOR CONTRIBUTIONS

VO and MS conceived and designed the study. AJurisic, CR and MS designed and performed the experiments and interpreted the data. BS and AJauch performed and interpreted M-FISH experiments. PT performed PUB-NChIP-seq experiments. LS performed bioinformatic analysis and interpretation of high throughput sequencing data. CSS, KE, ML and MS directed and critically revised the article. AJurisic, MS and VO wrote the manuscript.

